# Value Encoding in the Globus Pallidus: fMRI reveals an interaction effect between reward and dopamine drive

**DOI:** 10.1101/231910

**Authors:** Vincenzo G. Fiore, Tobias Nolte, Francesco Rigoli, Peter Smittenaar, Xiaosi Gu, Raymond J. Dolan

**Affiliations:** School of Behavioral and Brain Sciences, University of Texas at Dallas, 2200 West Mockingbird Lane, Dallas, TX 75235, USA; Wellcome Trust Centre for Neuroimaging, University College London, 12 Queen Square, London WC1N 3BG, UK; Max Planck UCL Centre for Computational Psychiatry and Ageing Research, 10-12 Russell Square, London, WC1B 5EH, United Kingdom

**Keywords:** Globus Pallidus, Indirect Pathway, Basal Ganglia, Dopamine, Parkinson’s disease

## Abstract

The external part of the globus pallidus (GPe) is a core nucleus of the basal ganglia (BG) whose activity is disrupted under conditions of low dopamine release, as in Parkinson’s disease. Current models assume decreased dopamine release in the dorsal striatum results in deactivation of dorsal GPe, which in turn affects motor expression via a regulatory effect on other nuclei of the BG. However, recent studies in healthy and pathological animal models have reported neural dynamics that do not match with this view of the GPe as a relay in the BG circuit. Thus, the computational role of the GPe in the BG is still to be determined. We previously proposed a neural model that revisits the functions of the nuclei of the BG, and this model predicts that GPe encodes values which are amplified under a condition of low striatal dopaminergic drive. To test this prediction, we used an fMRI paradigm involving a within-subject placebo-controlled design, using the dopamine antagonist risperidone, wherein healthy volunteers performed a motor selection and maintenance task under low and high reward conditions. ROI-based fMRI analysis revealed an interaction between reward and dopamine drive manipulations, with increased BOLD activity in GPe in a high compared to low reward condition, and under risperidone compared to placebo. These results confirm the core prediction of our computational model, and provide a new perspective on neural dynamics in the BG and their effects on motor selection and motor disorders.

## 1. Introduction

Pioneering studies (Albin et al., 1991; Albin et al., 1989; Alexander et al., 1986; DeLong, 1983, 1990; Smith et al., 1998) investigating the function of the basal ganglia (BG) proposed these interconnected nuclei play a fundamental role in action facilitation, and in the regulation of voluntary movement. Subsequent local connectome analyses resulted in further model developments (Frank, 2005, 2006; Gurney et al., 2004; Humphries et al., 2006; Nambu, 2004), including the suggestion that biophysical dysfunctions in the BG circuit might explain specific behavioural disorders and diseases (Obeso et al., 2014). These models propose that the output of the BG exerts a tonic inhibition of all motor commands to mediate a *gating* function. This output activity combines information conveyed through several converging pathways, termed direct, indirect and hyperdirect. It is hypothesised that these pathways compete to control activity of BG output nuclei, resulting in general inhibition or selective disinhibition (Calabresi et al., 2014; Nelson and Kreitzer, 2014; Schroll and Hamker, 2013). The direct pathway is thought responsible for motor facilitation and selective disinhibition and conveys cortical information via striatal medium spiny neurons rich in D1 receptors, to the output nuclei of substantia nigra pars reticulata (SNr) and globus pallidus pars interna (GPi). The indirect pathway conveys cortical information via D2-enriched spiny neurons in the striatum to the globus pallidus pars externa (GPe), which has efferent connections towards SNr, GPi, sub-thalamic nucleus (STN), striatum and parafascicular thalamic nucleus (Abdi et al., 2015; Gittis et al., 2014; Mastro et al., 2014). Finally, the hyperdirect pathway bypasses the striatum as input structure, and conveys cortical information to the SNr and GPi via the STN, establishing a recurrent circuit with the GPe (Smith et al., 1998).

A classic view states that striatal dopamine (DA) release modulates activity of the internal nuclei of BG, arbitrating the competition among the converging pathways for the control of the output of the system (Albin et al., 1991; Albin et al., 1989; DeLong, 1990; Frank, 2011). It is generally assumed that low DA drive in the dorsal striatum, such as seen in Parkinson’s disease (Rodriguez-Oroz et al., 2009), results in an increased signalling in D2-enriched striatum, which in turn causes decreased activity in the GPe (Chan et al., 2011; Filion and Tremblay, 1991). In a cascade effect, the suppression of activity in the GPe enhances activity in the STN, and in the BG output nuclei SNr and GPi, to inhibit motor expression. This model establishes a quasi-linear relationship between striatal DA release, pathway activity and behaviour (e.g. see: Frank et al., 2007; Nambu, 2004). Low DA activity is associated with increased indirect pathway signalling and motor suppression, while high DA activity is associated with increased direct pathway activity and motoric facilitation (Obeso et al., 2008a; Obeso et al., 2008b).

However, recent studies provide evidence that conflicts with both an hypothesis of competing pathways and an assumption of a linear correlation between striatal DA release and neural activity or resultant behaviour (Calabresi et al., 2014; Nambu, 2008; Nelson and Kreitzer, 2014). Firstly, concurrent activity in direct and indirect pathways has been found during motor initiation, highlighting the role played by the indirect pathway in triggering contraversive movements (Tecuapetla et al., 2016; Tecuapetla et al., 2014). Secondly, subthalamic deep brain stimulation ameliorates Parkinson’s disease motor symptoms by overactivating STN, whose signalling is already enhanced by low DA release (Fiore et al., 2016; Galati et al., 2006; Hashimoto et al., 2003). Finally, the role played by the GPe in the indirect pathway has been revisited (Gittis et al., 2014), disputing an assumption that this nucleus functions as a relay between D2-striatum and STN. In fact, the GPe is now seen as composed of heterogeneous neural populations (Abdi et al., 2015; Mallet et al., 2012; Mastro et al., 2014; Mastro and Gittis, 2015; Saga et al., 2017) that express complex patterns of activity (Bevan et al., 2002; Brown, 2007; Chiken and Nambu, 2016; Mallet et al., 2008a; Mallet et al., 2008b), suggesting its computational role in the BG also needs to be revisited.

Using a neural model, we have recently proposed a different perspective on BG function and neural dynamics, with particular attention on the role played by the GPe and the indirect pathway. This theory assumes that cortical information in direct and indirect pathways is either compressed or amplified as a function of striatal DA release. The transformed information in the BG pathways is integrated in the output nuclei and propagated back to its origin in the cortex. Due to this recurrent circuit, information enhancement in the direct pathway results in increased circuit gain, strong attractors and stable state transitions (Fiore et al., 2014), whereas information compression in the indirect pathway results in decreased circuit gain, shallow attractors and meta-stable dynamics (Hauser et al., 2016) or even oscillations (Fiore et al., 2016). This change of perspective is essential to account for both the classic DA-related coarse arbitration between suppression and facilitation of motor and cognitive functions, as well as the new data suggesting a more complex and finely grained regulatory role of the indirect pathway (Gittis et al., 2014; Tecuapetla et al., 2016; Tecuapetla et al., 2014; Vicente et al., 2016). In particular, our simulations show the encoding of the cortical input in the internal nuclei of the BG interacts with the striatal DA release, resulting in non-linear dynamics and in broadening the set of the possible functions and dysfunctions associated with BG gating. Crucially, our model predicts that striatal DA modulation results in information compression in the GPe, under basal or high DA drive, and information amplification in the GPe under reduced DA drive (**Fig. 1**). A similar interaction, but with the opposite direction of information compression and amplification is predicted for the BG output nuclei (**Fig. 1**).

**Figure 1.**
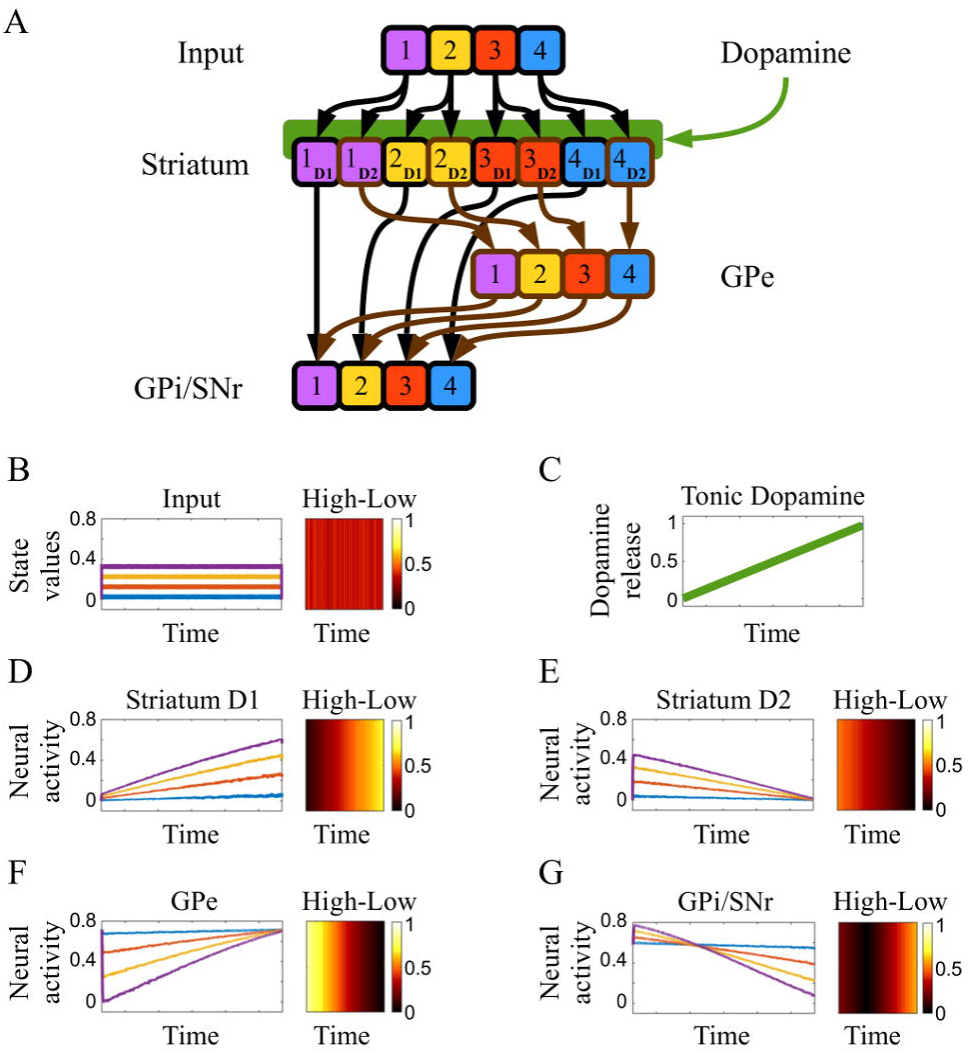
Neural model and simulated neural dynamics. A) Schematic representation of the architecture used to represent a simplified BG circuit and simulate the activity of its interconnected nodes over time. Four simulated sensorimotor or action-state values (B), putatively encoded in cortical signals reaching the striatum, are constant during the whole simulation. Tonic DA release targeting the striatum (C) slowly increases during the simulation, affecting the way the input is encoded in the striatum. Striatal D1-enriched units (D) amplify the differences among the inputs as a direct function of DA release. Conversely, striatal D2-enriched units (E) show the higher differentiation under low DA release, as the input signals are compressed towards the end of the simulated time, in association with high DA release. GPe (F) and GPi/SNr (G) receive the input after it is processed by D2 and D1 enriched striatum, respectively. Due to the inhibitory afferent connections, the GPe mirrors the signal received from the D2 striatum. Finally, GPi/SNr receive conflicting inhibitory information from D1 striatum (direct pathway) and GPe (indirect pathway), resulting in the compression of signal differences, at low dopamine release. The key prediction of this computational hypothesis is further illustrated in the heatmaps for each BG nucleus, where we represent the differences in simulated neural activity between the encoding of high and low action-state values. This difference is tested in our within-subject fMRI paradigm, as the model predicted increased High-Low differentiation in the GPe under a condition characterised by low dopaminergic release, in comparison with basal or high dopaminergic conditions.

To test our predictions, we examined twenty-four healthy volunteers performing a reward-based motor task during fMRI. We used a within-subject comparison of behaviour and fMRI blood-oxygen-level dependent (BOLD) activity in a 2x2 design where we manipulated DA drive and perceived action-state values. This design was used to test the predicted interaction effect, as we expected to find enhanced action-state value differentiation in the GPe under a low DA drive condition, induced by treatment with risperidone.

## 2. Materials and Methods

### 2.1 Neural Model

We used a simplified neural architecture (**Fig. 1A**) to illustrate how DA release dynamically interact with action-state value representations in the striatum, GPe, GPi and SNr. We assumed four inputs, kept constant through the simulation (**Fig. 1B**), and representing the cortical encoding of the values associated with actions performed under specific environmental conditions, hence action-state values (cf. Sarsa algorithms as introduced in: Rummery and Niranjan, 1994). These inputs are propagated via parallel connectivity towards a layer of neural units representing the striatum. During the same time interval, a simulated release of tonic DA gradually increases (**Fig. 1C**) as it reaches the striatal units. These units respond in a different way to the incoming inputs and DA, depending on the presence of either D1 and D2 receptors (**Fig. 1D,E**). The resulting activity reaches via inhibitory connections the internal nucleus GPe (**Fig. 1F**) and the output nuclei GPi and SNr (**Fig. 1G**), which are also affected by the GPe, via inhibitory connections. For illustrative purpose, this BG circuit has been simplified by considering the output nuclei as identical (hence labelled GPi/SNr, **Fig 1G**) and by using a baseline positive activity in place of the excluded excitatory signal derived by the sub-thalamic nucleus (for a more detailed version of the model, see: Fiore et al., 2016; Hauser et al., 2016). The activity of all units in this model is described in a continuous time differential equation (1) a positive saturation transfer function (2):

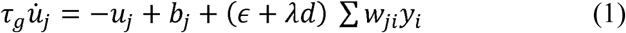

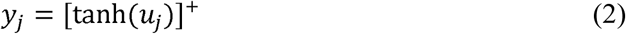

where the action potential of a unit *u_j_* is updated at a pace determined by the time constant *τ_g_*, and depending on the value of a baseline *b_j_* (constant per nucleus) and an input Σ*w_ji_y_i_*, weighed by (*∈* + *λd*). The two constants *∈* and *λ* respectively regulate the amount of input signal that is independent of the presence of DA (*d* in equation 1), and the response of the DA receptor. In the simulated striatum, *λ*=2 and *λ*=-1.5 simulate D1 and D2 receptors respectively, whereas *λ*=0 for all other units. The model was developed using Matlab.

### 2.2 Participants

24 healthy volunteers (17 females, 23 right handed), age 25.1±0.9, weight 60.4±7.3 kg, were recruited for this study via an advertised mailing list hosted by the Institute of Cognitive Neuroscience at University College London. Selection criteria included weight (inclusion range: 50kg to 70kg) and age (inclusion range of 20-40 years). These criteria were based on a previous study (Fiore et al., 2016) to enhance chances of a consistent effect of the administered DA antagonist across subjects. Participants taking any medication, or with a history of mental disorder or drug abuse were excluded from the study. Subjects were asked to avoid consumption of alcohol, coffee, tea, energy drink, or any similar stimulant 12 hours prior to each session, but they were not assessed with a toxicology test. Informed consent was obtained from all participants, who were made aware that they could quit the study at any time. The ethics committee of the University College London approved the experiment. Data collected in 4 participants (3 females) had to be discarded due to: (n=2) malfunctioning of the apparatus for recording behavioural responses (see subsection 2.3), (n=1) incomplete data due to drop out, and (n=1) incorrect positioning of the participant with respect of the coiler in one of the two sessions.

### 2.3 Experimental Design and Statistical Analysis

The study was designed to define 2x2 conditions, with reward (high vs low) and DA modulation (placebo vs DA antagonist risperidone) as principal variables. All participants were requested to participate in two sessions: a placebo condition and a second involving DA manipulation, where we administered 0.5 mg of the DA antagonist risperidone. This is an antipsychotic medication, mainly used to treat schizophrenia and selected in the present study on account of its binding affinity with D2 (3.57 Ki [nM], antagonist) and D1 receptors (244 Ki [nM], antagonist). Like other DA antagonists (e.g. see: asenapine, blonanserin, clozapine, olanzapine, zotepine), risperidone also interacts with serotonin (5-HT2) and noradrenaline (α1/2) receptor subtypes. Session order was counterbalanced across subjects and the pharmacological manipulation followed a double blind procedure. An authorised medical doctor (T.N.) was present during the study and administered a glass of juice 45 minutes to one hour prior to the start of the task in the scanner. The DA antagonist was dispersed in juice in half the cases. Dosage and schedule, tested in a previous study (Fiore et al., 2016), was conceived to reduce the incidence of extrapyramidal side effects, whilst testing the volunteers when the drug was at its maximum effect (mean peak plasma concentrations of risperidone occurs at about 1 hour). An interval of at least 7 days between sessions has been applied.

Each session consisted of three blocks, lasting 10 to 12 minutes each. Each block involved 72 trials, organised as follows: 2.5-3.5 seconds for the fixation cross, 1.5 second for the reward condition and 2-4 seconds for the action response (**Fig. 2**). Trial order was pseudorandomized to alternate between reward (either 10 points or 1 point per second spent applying the correct force) and action type. A calibration phase and a short training were required prior to each session. During calibration participants were asked to apply a force that they could feel comfortable with for the duration of the entire task. After the training (150 seconds circa per 24 trials), the participants were allowed to repeat the calibration phase if they wished so, but they could not change these settings after the beginning of the first block of trials. This procedure also allowed the participants to establish a motor memory for the force to be applied, avoiding learning processes during the actual task.

**Figure 2.**
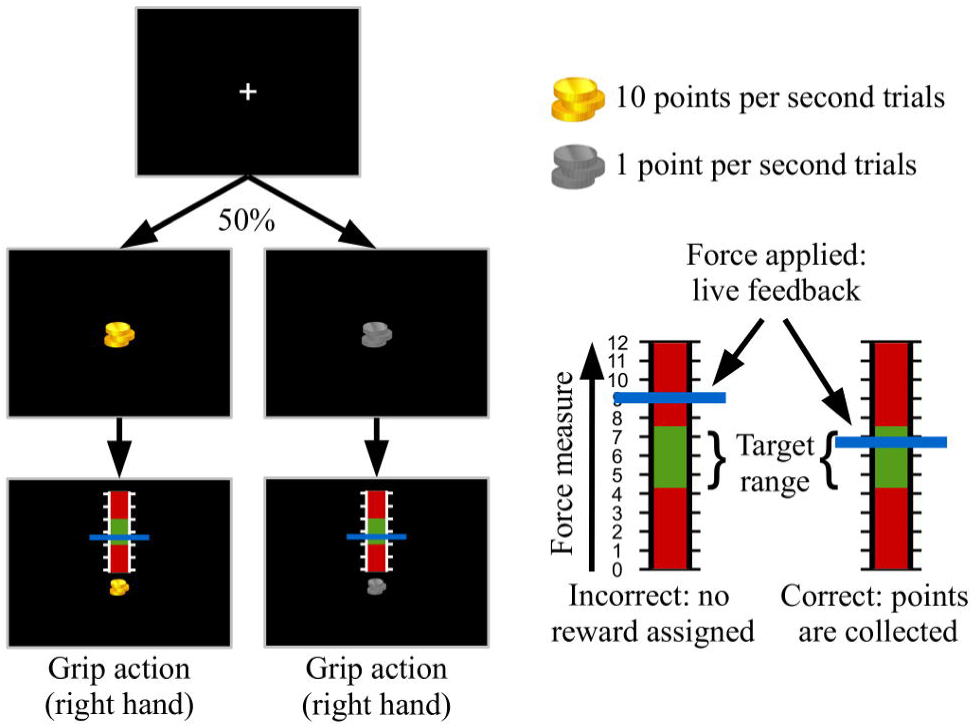
Schematic representation of the computer task. fixation cross is followed by a symbol indicating the starting trial is characterised by either high reward (“gold coins”) or low reward (“iron coins”), with equal probabilities. Each condition is followed by the motor part of the task, where a red and green scale-like image is presented to the participant on a black screen. This indicates the moment the “grip action” has to be performed, applying a sustained force on the apparatus, with the right hand, until the end of the trial, for 2-4 seconds. The force applied by a participant is constantly recorded and reported on screen by means of a horizontal blue line that moves vertically on the scale. The participants collect points proportionally to the reward of the trial, per each second spent applying the correct force, i.e. having the feedback blue line positioned in the “green zone” of the scale.

The task required participants to apply a force established during the calibration phase so as to score points and maximise their reward. Either 10 points or 1 points per second were assigned, depending on the reward condition, proportional to the time spent maintaining a blue bar representing a live feedback (graphical update: 0.1 seconds) of the force applied, in a “green zone”. This target area was marked at the centre of an illustrative force meter that appeared on the screen at the beginning of each trial (**Fig. 2**). The force required to reach the centre of this “green zone” was set during the calibration. This procedure was conceived as an incentive to facilitate participants to always apply the same force, independently of the reward associated with the trial, therefore avoiding force-related differences in BOLD activity (Pessiglione et al., 2007; Spraker et al., 2007; Vaillancourt et al., 2004). Participants were aware that the compensation was computed as £1 for every 125 points and were reminded they could score up to 40 points per high reward trial, or 4 points per low reward trial.

The experiment was replicated twice for two motor actions. In one context participants were required to use the right hand on a hand grip apparatus equipped with pressure sensors. In the second case the sensors were located on a pedal activated with the right foot. Unfortunately, due to equipment failure, data recorded with the pedal had to be discarded in the present analysis. More precisely, to the position of the participants in the scanner (the leg had to be approximately 45 degrees above the pedal) and the fact that the apparatus itself could be pushed away a few centimetres during the trials, resulted in the pressure sensors in the pedal to often confound changes of weight on the foot on the pedal as voluntary motor actions.

In the analysis of both behaviour and fMRI BOLD activity, repeated measures ANOVA was used to test the presence of an interaction effect between the two variables of reward and DA manipulation. Behavioural analysis focused on the measure of reaction time, calculated as the time required to apply the rewarded force. The time counter started when a visual cue for the motor action appeared on the screen and it was stopped when the participants applied a force within the required range (i.e. the force feedback system signals the participants that the “green zone” has been reached). Repeated measures ANOVA were used to test the presence of an interaction effect when considering beta values extracted from ten independent regions of interests (ROI) under the four conditions defined by the 2x2 design.

### 2.4 fMRI data acquisition and preprocessing

A 3-dimensional sequence with a resolution of 2 mm was chosen for a 3-Tesla MR scanner (Siemens) at the Functional Imaging Laboratory, Wellcome Trust Centre for Neuroimaging at UCL. Structural imaging was carried out with a resolution of 1 mm, Multi Parametric Maps. Functional images were acquired with a 3D echo-planar imaging (EPI), flip angle = 0°, volume repetition time of 3.5 seconds, echo time of 30 ms, 52 slices, Matrix size 96 × 108, echo spacing of 0.78 ms, transverse orientation, and a resolution of 2.0x2.0x2.0 mm. FMRI data preprocessing was performed using statistical parametric mapping (SPM12, Wellcome Department of Imaging Neuroscience). The functional scans were realigned to the first volume, coregistered to the T1 image, and normalized to a standard MNI (Montreal Neurological Institute) template. The scans were only spatially smoothed for the whole brain analysis (=6 mm), but they were not spatially smoothed for the ROI analysis, due to the small volumes of the chosen ROIs.

### 2.5 General linear modelling (GLM) of fMRI data

Event-related analyses of the fMRI data were conducted using statistical parametric mapping (SPM12; Wellcome Department of Imaging Neuroscience, London, UK). GLM (Friston et al., 1994) was conducted for the functional scans from each participant by modelling the observed event-related BOLD signals and regressors to identify the relationship between the task events and the hemodynamic response. Regressors related to all events were created by convolving a train of delta functions representing the sequence of individual events with the default SPM basis function, which consists of a synthetic hemodynamic response function composed of two gamma functions (Friston et al., 1998; Friston et al., 1994). We combined both sessions and concatenated the six total runs to include in one model the regressors of all 4 conditions from the 2 (drug: placebo [pla] vs. risperidone [ris]) by 2 (reward: high reward [HR] vs. low reward [LR]) design: pla-LR, pla-HR, ris-LR, ris-HR. For the whole brain analysis we time-locked events to the moment of presentation of the reward symbol, with a duration of zero, so as to test the main effects of the two variables, independently of the actions performed. For this analysis, we used a combined threshold of p<.005 with a 50 voxel extent to highlight significant differences (e.g. see: Lieberman and Cunningham, 2009). For the ROI analysis, we meant to measure BOLD activity in association with the sustained motor response of each trial. Thus, the events were time-locked to a time interval having the visual cue signalling the beginning of motor action as a start and the end of each trial as the end of the event. Linear contrasts of the parameter estimates were implemented to identify effects in a within subject comparison.

### 2.6 Regions of Interest

To test our key hypothesis and the predictions of the model, we focused on the internal and output nuclei of the BG. Participants were instructed to perform their actions with the right hand, therefore we expected to find the key changes in the encoding of action-state values in BOLD activity of the left hemisphere, localised in the areas associated with motor execution. The dorsal segment of both GPe and GPi is responsible for encoding sensorimotor command selection and execution (Draganski et al., 2008; Pessiglione et al., 2007; Saga et al., 2017), therefore we constructed different masks to allow separate analysis for dorsal and ventral areas of both GPe and GPi, which were analysed bilaterally for comparison. Finally, we analysed bilateral activation of SNr, which represents the output nucleus processing mainly information, such as expected future outcomes, encoded in the ventral cortico-striatal loop (Draganski et al., 2008; Haber, 2003; Jahanshahi et al., 2015).

The masks for all ROIs were manually defined to account for the dorsal/ventral separation and to focus only on the SNr, thus excluding the adjacent dopaminergic area of the SN pars compacta. As a reference for the nuclei in their entirety, we used recently published and freely available online atlases of the BG (Smittenaar et al., 2017; Xiao et al., 2012; Xiao et al., 2015; Xiao et al., 2017). The sensorimotor section of both GPe and GPi accounts for roughly half of their entire volume (Rodriguez-Oroz et al., 2009; Romanelli et al., 2005) and the two parts of the GP extend on the horizontal plane roughly between the values y=-14 and y=7 (GPe) and y=-11 and y=3 (GPi). Thus we divided dorsal and ventral segments on the horizontal plane at the value y=-3. Within the nuclei of interest, we used the ROI to extract signal under the four conditions of the experimental design (pla-LR, pla-HR, ris-LR, ris-HR). All ROIs were defined using the software MRIcron and MarsBaR, used jointly with SPM 12.

## 3. Results

### 3.1 Simulations and predictions

The neural model illustrates how the nuclei of this simplified BG circuit process and encode a constant four dimensional input, putatively representing four action-state values perceived by the agent. The difference between high and low values reaching the layer simulating the striatum is kept constant through. Nonetheless, the representation of these inputs and the difference between highest and lowest values is either amplified or compressed, as a function of DA release (**Fig 1**). The simulations highlight increased value differentiation in the GPe, under low dopaminergic conditions (**Fig 1G**), and in the GPi or SNr, under high dopaminergic condition (**Fig 1F**). Normally, this process of amplification and compression of information in the direct and indirect pathway interacts with the recurrent connectivity of the cortico-thalamo-striatal circuits, resulting the generation and modulation of the attractor states in the system (Fiore et al., 2016; Fiore et al., 2014; Hauser et al., 2016). In the present model, we limit our simulations to a feed-forward neural network to simplify the representation of these neural dynamics. Finally, due to the design of our DA manipulation, in the present study we focus on the condition of low DA drive, establishing a comparison with baseline condition.

### 3.1 Behavioural results

We used repeated measures ANOVA to test the effects reward and DA manipulation have on the measure of reaction time (**Fig 3**). No main effect was found for any of the two variables (DA antagonist vs placebo: F=2.75, p=.11; high vs low reward: F=1.85 p=.19) with no interaction effect (DA*Reward, F=.31 p=0.58). The behavioural analysis highlighted the presence of an outlier (participant 5, **Fig. 3**) who was consistently slower than the remaining participants under the condition of risperidone. The slow responses, jointly with the limited time (2-4 seconds) available to produce an action, led this participant to miss several trials (a total of 17 trials against an average of 1.18 for the remaining 19 participants). We will consequently report significant ROI results with and without this participant.

**Figure 3.**
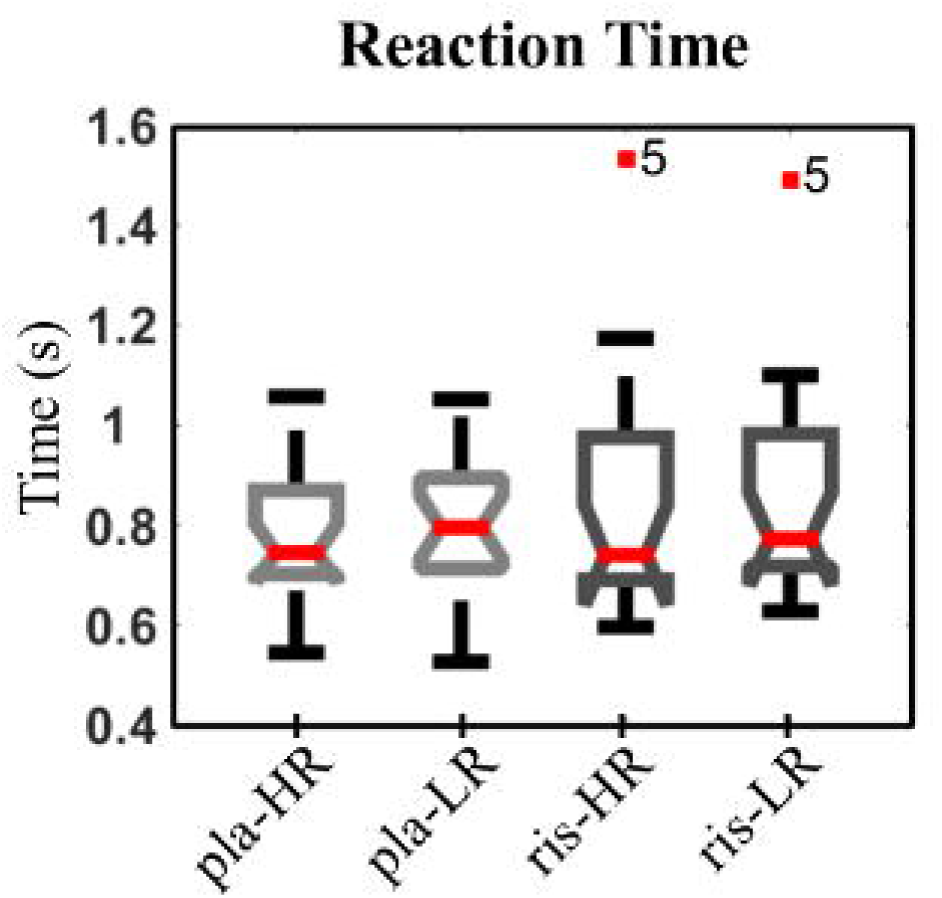
Boxplot representation of the distribution of reaction times under the conditions of high and low reward (HR and LR, respectively), combined with the DA drive condition of placebo (pla) or risperidone (ris). Participant 5 is highlighted as outlier in terms of RTs recorded under the condition of risperidone.

### 3.2 fMRI results

We measured the whole brain response to the manipulation of each variable, testing BOLD signal when contrasting either placebo vs risperidone conditions or high vs low reward conditions. By comparison with the risperidone condition, the placebo condition was found to be associated with greater BOLD response in orbitofrontal cortex (right hemisphere) and dorsal striatum (bilateral, p<.005, voxel extent: 50, main peaks of activity: 22,10,-6; 28,22,-8, and −20,8,−8, with right hemisphere results surviving whole brain correction: p_FWE_-corr__<.001; **Fig. 4A**). The contrast between high reward and low reward conditions showed the former was associated with an increase in BOLD response in the pre-frontal cortex (p<.005, voxel extent: 50, peak of activity: −2,56,10, which does not survive whole brain correction; **Fig. 4B**).

**Figure 4.**
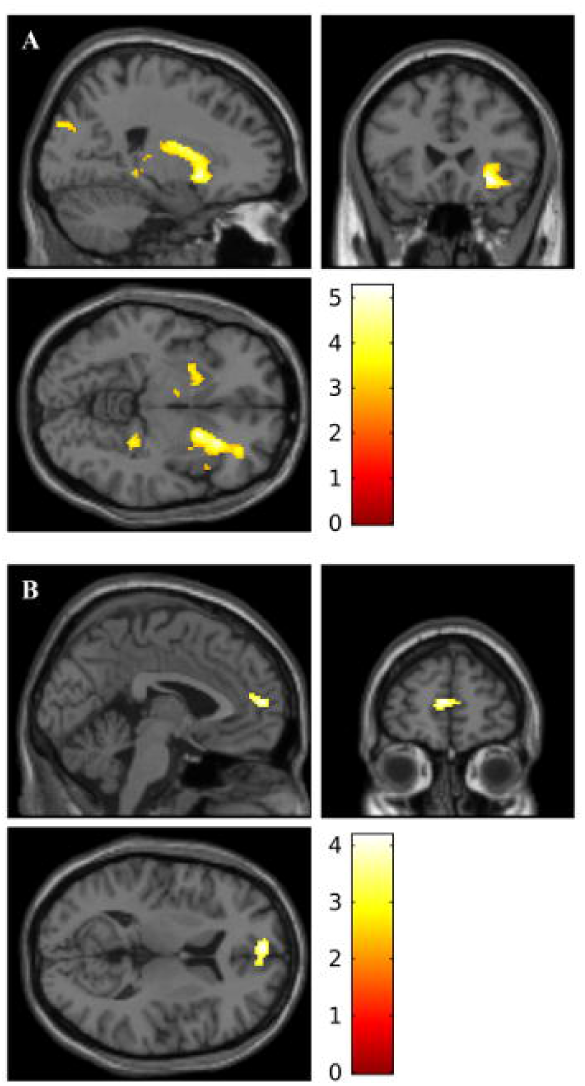
Whole brain activity reported with a threshold of p < .005 and a 50 voxel extent. (A) The first contrast (coordinates for the image: 22, 24, −6), between placebo and risperidone condition, shows BOLD activity in the putamen (bilateral), caudate (right hemisphere) and orbitofrontal cortex (right hemisphere). (B) The second contrast (coordinates for the image: - 2, 56, 10), between high and low reward presentation, reveals BOLD activity in the prefrontal cortex.

For the ROI analysis, the extracted signals under the four conditions of the experimental design have been used to run 2 by 2 repeated measures ANOVA for each ROI. We found no main effect for reward manipulation when considering the selected ROIs in GPe (left dorsal: F=.002, p=.96, left ventral: F=1.59, p=.22, right dorsal: F=.12, p=.73, right ventral: F=2.11, p=.16), GPi (left dorsal: F=1.59, p=.22, left ventral F=1.29, p=.27, right dorsal: F=.97, p=.34, right ventral: F=.07, p=.80) and SNr (left: F=.10, p=.75, right: F=.21, p=.66). Similarly, we found no significant main effect in GPe (left dorsal: F=1.27, p=.27, left ventral: F=.002, p=.97, right dorsal: F=2.54, p=.13, right ventral: F=.72, p=.41), GPi (left dorsal: F=.02, p=.90, left ventral: F=1.02, p=.33, right dorsal: F=.004, p=.95, right ventral: F=2.29, p=.15) and SNr ( left: F=.68, p=.42, right: F=.01, p=.92), when considering the drug manipulation.

Our core hypothesis was a predicted interaction effect in the dorsal GPe, left hemisphere, involving increased signal in presence of high values in comparison to low values, under reduced dopaminergic drive (cf. **Fig. 1F**). The results support this core prediction, as repeated measures ANOVA revealed a significant interaction effect of the two variables in the GPe (left dorsal: F=6.53, p=.02; if the behavioural outlier is included: F=3.52, p=.076). This interaction effect was limited to the area responsible for motor execution, and was not found in any of the remaining GPe ROIs (left ventral: F=.00, p=.99, right dorsal: F=.17, p=.68, right ventral: F=.00, p=.99). As predicted, extracted beta from dorsal GPe ROI show the representation of the action-state values in the task changed direction. Under the placebo condition, BOLD activity was inversely correlated with reward, as highlighted by the mean within subjects difference between extracted beta values under HR and LR conditions (mean differences HR-LR: −0.13, **Fig.5**). Conversely, under risperidone condition, GPe increased its BOLD signal in association with high reward (mean difference HR-LR: +0.14, **Fig.5**).

**Figure 5.**
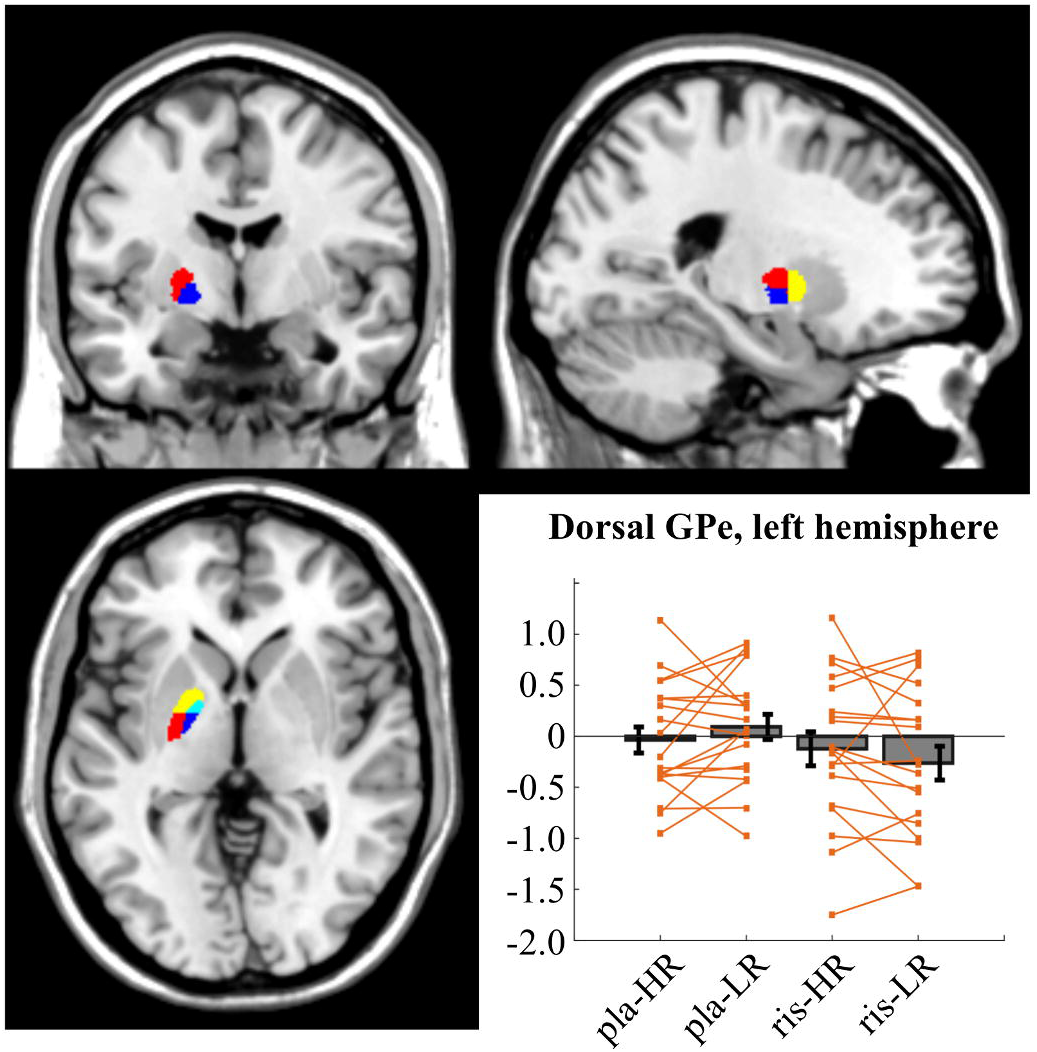
Illustration of the maps used for the globus pallidus ROIs in the left hemisphere (coordinates for the image: −22, −4, 0) and extracted beta values for the left dorsal GPe. In the map, dorsal and ventral GPe are highlighted in red and yellow, respectively, whereas dorsal and ventral GPi are highlighted in blue and cyan, respectively. Extracted values are reported as bars with mean and standard error for the four conditions characterising the experimental design: high vs low reward (HR - LR) and placebo vs risperidone (pla - ris). Single data points are reported (orange) per each condition, linking values extracted under HR and LR conditions. Repeated measures ANOVA shows a significant interaction effect (F=6.53, p=.02), as the mean within subject difference changes from HR-LR=-0.13, under risperidone condition, to HR-LR=+0.14, under placebo condition. No main effect is reported for either variable. The beta values are reported after the exclusion of the behavioural outlier (participant 5).

Finally, we analysed the activity in the output nuclei of the BG, testing for the presence of the opposite interaction effect (cf. **Fig. 1G**). No significant effect was found in the GPi ROIs (left dorsal: F=.40, p=.53, left ventral: F=.18, p=.67, right dorsal: F=1.6, p=.22, right ventral: F=.21, p=.65). However, we found a significant interaction effect when analysing beta extracted from the right SNr (F=4.85, p=.04; if the behavioural outlier is included: F=5.32, p=.03), where participants 7 and 11 were outliers under one out of the four conditions (ris-HR and ris-LR, respectively). No interaction effect was found in the left SNr (F=.24, p=.63). The analysis shows the presence of a canonical response for reward encoding in the SNr under placebo condition, where BOLD activity directly mediates the presence of the expected outcomes (mean within-subject difference of extracted beta values: HR-LR= +0.21, **Fig. 6**). This representation is inverted under risperidone condition, as BOLD activity in the SNr shows an inverse correlation with the reward associated with the trial (mean difference HR-LR: −0.13 for HR and LR, respectively, **Fig. 6**).

**Figure 6.**
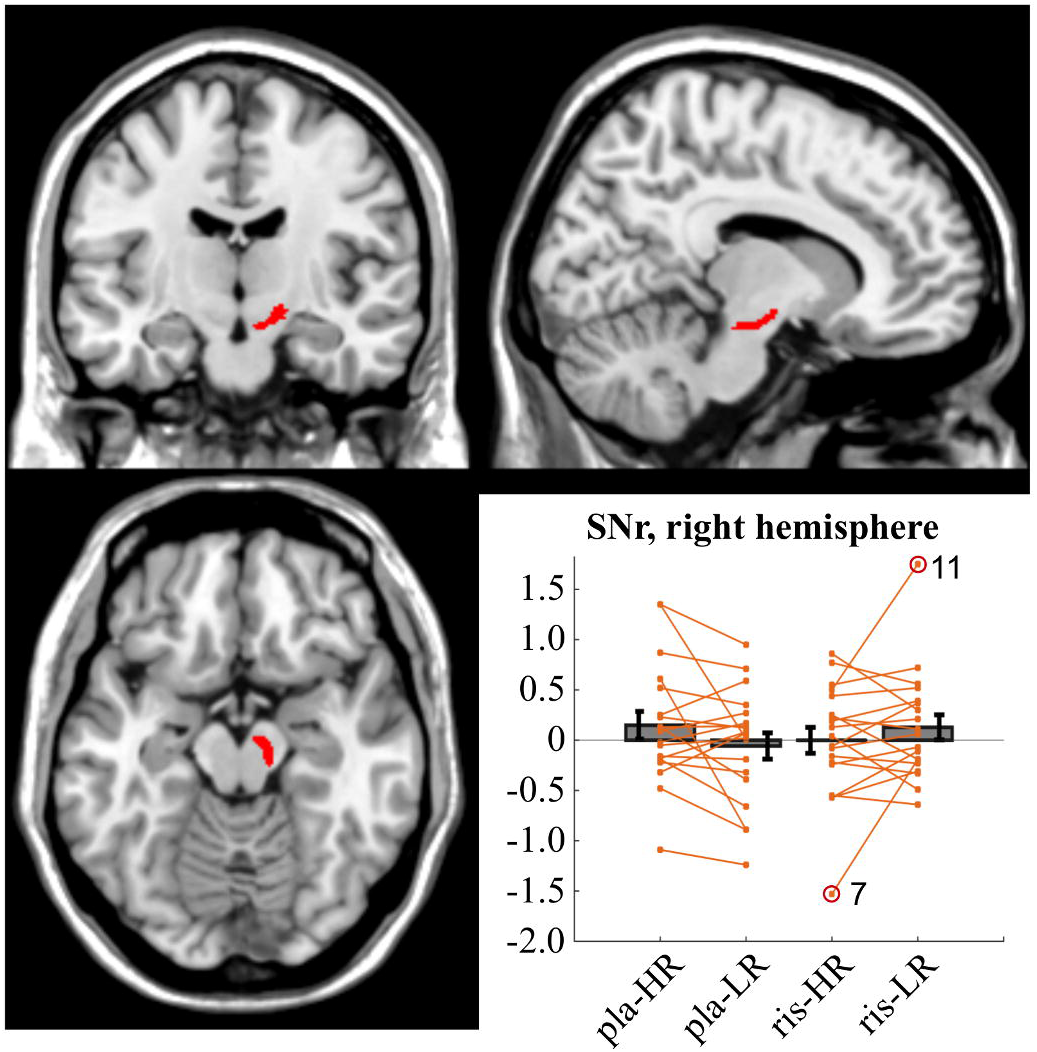
Illustration of the map used for the SNr ROI, right hemisphere (coordinates for the image: 11, −13, −15), and extracted beta values for this mask under the four conditions characterising the experimental design: high vs low reward (HR - LR) and placebo vs risperidone (pla - ris). Bars with mean and standard error are overlaid with single data points (orange) per each condition, where we have linked values extracted under HR and LR conditions. A significant interaction effect was found in the right SNr (F=4.85, p=.04), which was caused by a change of direction in the mean of the within subject difference: HR-LR=+0.21, under risperidone condition, HR-LR=-0.13, under placebo condition. No main effect is reported for either variable. The beta values are reported after the exclusion of the behavioural outlier (participant 5). Two further outliers for the beta values (participants 7 and 11) are also marked with a red circle.

## 4. Discussion

Classic interpretation of the BG function predicts that, under the condition of low DA drive, increased signalling in the indirect pathway takes the lead in the competition for the control of the output of the BG, resulting in motor suppression. Conversely, under high DA drive, increased signalling in the direct pathway results in motor facilitation (Albin et al., 1989;Alexander et al., 1986; DeLong, 1990; Smith et al., 1998). This hypothesis assumes that a specific role is played by the internal nuclei of the BG. Most significantly: 1) low DA drive and motor suppression are associated with reduced GPe activity and increased firing in the output nuclei of GPi and SNr; and 2) high DA drive and motor facilitation are associated with increased GPe activity and decreased firing in the output nuclei. This linear mechanistic explanation can account for a wide variety of motor and cognitive findings and dysfunctions, such as those related to DA deficiency in Parkinson’s disease (Obeso et al., 2014; Rodriguez-Oroz et al., 2009). Nonetheless, shortcomings of this theory have emerged in new findings that reveal a complex circuitry and activity patterns involving the GPe (Bevan et al., 2002; Gittis et al., 2014; Mallet et al., 2008a), as well as a previously unknown active role played by the indirect pathway in promoting specific motor activity (Nelson and Kreitzer, 2014; Tecuapetla et al., 2016; Tecuapetla et al., 2014). These findings led us to formulate a new, more comprehensive theoretical framework (Fiore et al., 2016; Hauser et al., 2016) which states that cortical information encoding context or state values and the related sensorimotor contingencies (Azzi et al., 2012; Montague et al., 2006; O’Doherty, 2014) can be either compressed or amplified in the BG pathways as a function of DA release (**Fig. 1**). This new interpretation implies activity in the GPe, GPi and SNr should be found to vary as a function of both DA drive and encoded action-state values.

In this study, we use a simplified neural architecture to illustrate the dynamics predicted in our model and we assess with fMRI the two competing predictions proposed by the classic interpretation and by our model of the BG, with a specific focus on the GPe. Namely, the former predicts a main effect of drug manipulation with stronger activity localised in the GPe under placebo condition. The latter predicts the presence of an interaction effect where the difference between the encoding of high and low action-state values is amplified under risperidone condition. To this end we used a motor task designed to manipulate two variables for a within-subject analysis. The DA drive was controlled by administering either placebo or DA antagonist risperidone. The action-state value was manipulated by explicit assignment of either low or high *expected rewards* to actions initiated, and sustained, for the entire duration of each trial. Our design involved a preliminary training phase and the task was then implemented without explicit trial-by-trial feedback, so as to avoid or reduce any learning during the experiment. This design allowed us to interpret the results in terms of how fixed action-state values (i.e. maximum theoretical rewards associated with a trial) are encoded in the BG nuclei, independent of learning and plasticity.

Our results show the two variables of DA drive and reward had expected main effects on striatum and orbitofrontal cortex. However, no main effect was found for the BOLD activity in any of the nuclei of the BG. In keeping with the prediction of our model, we found the interaction between the two variables had a significant effect: 1) in the left dorsal GPe, as the representation of action-state values improves under reduced DA drive; 2) in the right SNr, where we found the value representation is inverted when comparing placebo and risperidone conditions. Importantly, the participants were rewarded when responding with stereotyped actions, as changes in motor activity (e.g. pace or intensity of motor responses) have been reported to cause differences in BOLD responses (Pessiglione et al., 2007; Spraker et al., 2007; Vaillancourt et al., 2004). The results show the ability of the participants to apply and maintain a constant force (within a limited range) did not vary under the different conditions. Therefore, the task successfully avoids possible confounds, enabling us to associate variations in BOLD activity with the experimental manipulation of the two variables of interest. The use of ROIs, which were manually defined for the target regions in the BG on the basis of previous maps (Smittenaar et al., 2017; Xiao et al., 2012; Xiao et al., 2015; Xiao et al., 2017), jointly with the analysis of spatially unsmoothed data, also prevented possible confounds derived from activity in adjacent areas. The interaction effect predicted in the GPe was found only in the dorsal segment, left hemisphere, thus in the area expected to encode state values associated with the execution of motor activity with the right hand (Draganski et al., 2008; Pessiglione et al., 2007; Saga et al., 2017). Finally, our model predicted the GPi would be found to express an interaction effect similar to the SNr, but no significant change in BOLD activity was found for the GPi. In keeping with the model, we hypothesise a condition characterised by higher than normal DA drive might help highlighting an effect in the GPi. Further investigations are required to test and possibly validate this hypothesis.

Despite important limitations, such as an unbalanced gender representation and a relatively small sample size, our findings nevertheless challenge the common view of the linear interaction between DA release and information processing in the BG. In particular, by highlighting that activity in GPe varies as a function of both DA drive and action-state values, our data offer some support for an alternative model of both motor and cognitive dysfunctions associated with disrupted striatal DA release. Further investigation is required to establish an unequivocal link between changes in activity in the GPe and selection switching or pattern generation, as has been suggested in recent work (Fiore et al., 2016; Tecuapetla et al., 2016; Tecuapetla et al., 2014; Vicente et al., 2016). This is particularly important when considering disorders associated with DA dysregulation, such as Parkinson’s disease. The possibility to generate increased motor switching, as well as the classic motor suppression, grants the model the required plasticity to simulate different behavioural phenotypes associated with the same DA biophysical dysfunction.

## 5. Conclusions

Our study provides insights into the way action-state values are encoded in the internal nuclei of the BG and the GPe in particular. Our results show that changes in mean field potentials in the GPe-suggested by the reported changes in BOLD activity- are not limited to DA manipulations, as it is often assumed. As predicted by our model, action-state values are encoded in the GPe and their differences are amplified under the condition of low dopaminergic release. We hypothesise this incorrect encoding interferes with the healthy selection process performed in the basal ganglia, under conditions such as Parkinson’s disease. A better understanding of BG dynamics can heavily impact the possibility to develop treatments for motor disorders, such as deep brain stimulation.

## Acknowledgements

The authors would like to thank Alphonso Reid for the help provided in setting up the response apparatus. The Wellcome Trust Centre for Neuroimaging is supported by core funding from the Wellcome Trust (091593/Z/10/Z). This work was supported by the Wellcome Trust (Ray Dolan Senior Investigator Award 098362/Z/12/Z). T.N. is supported by the Wellcome Trust Principal Research Fellowship to Professor Read Montague. X.G. is supported by a faculty start-up grant from UT Dallas and the Dallas Foundation.

